# Gut barrier integrity biomarkers are associated with increased inflammation and predict disease status in hospitalized COVID-19 patients

**DOI:** 10.1101/2025.03.28.645914

**Authors:** Christopher M. Basting, Ty A. Schroeder, Kathie G. Ferbas, Robin R. Shields-Cutler, Nicole H. Tobin, Adrian Velez, Erik Swanson, Courtney A. Broedlow, Robert Langat, Luca Schifanella, Carolyn T. Bramante, Grace M. Aldrovandi, Anne Rimoin, Otto O. Yang, Jennifer A. Fulcher, Nichole R. Klatt

## Abstract

The COVID-19 global pandemic persists as an endemic disease with frequent case spikes and a significant continued burden on public health. Although most COVID-19 cases are asymptomatic or mild, severe infections requiring hospitalization have resulted in likely more than 14 million cumulative deaths to date. One hallmark of severe COVID-19 is a dysregulated immune response that leads to systemic inflammation and contributes to disease severity and mortality but is not explained by viral replication alone. Severe COVID-19 has been shown to disrupt the gut microbiome and increase intestinal permeability which may contribute to immune dysregulation and systemic inflammation. In this study, we investigated the differences in plasma biomarkers for microbial translocation and gut barrier damage as well as circulating cytokines between healthy volunteers and patients hospitalized with COVID-19. We then performed a correlation analysis to understand how the relationships between these plasma biomarkers differed and used a random forest model to assess their accuracy in distinguishing between these two groups. Our results demonstrated that hospitalized COVID-19 patients have elevated concentrations of pro-inflammatory cytokines and markers of microbial translocation, and that the relationships between these biomarkers were significantly altered compared to healthy volunteers, especially those related to the mucosal associated homeostatic cytokines IL-17A and IL-23. Furthermore, IL-6 and LBP were the top biomarkers for prediction accuracy in our random forest model, highlighting the importance of managing microbial translocation in COVID-19 and its potential utility as a biomarker for disease severity.

**IMPORTANCE:** COVID-19 continues to be a burden on public health. Understanding how plasma biomarkers differ between healthy and acutely infected individuals can help in understanding pathogenesis and predict disease severity. It has been demonstrated that COVID-19 disrupts the gut microbiome and intestinal permeability, which contributes to systemic inflammation. Our study highlights the link between gut barrier integrity and inflammation in hospitalized SARS-CoV-2 cases, demonstrating the usefulness of gut barrier integrity biomarkers in predicting disease severity, and offers insights for therapeutic interventions in acute COVID-19 infection.

## INTRODUCTION

The SARS-CoV-2 virus is responsible for Coronavirus Disease 2019 (COVID-19) which continues to threaten public health as an endemic disease with intermittent acute spikes in incidence and persistent post-acute sequelae^1^. Viral infection is facilitated through binding of the viral spike protein to angiotensin converting enzyme 2 receptor (ACE2), a widely expressed receptor found throughout the body but especially in the lungs and gastrointestinal (GI) tract^2^. This widespread expression allows SARS-CoV-2 to infect multiple organ systems^3^, which may explain why COVID-19 is often accompanied by involvement beyond pulmonary symptoms, such as those arising from the GI tract^4–6^.

A hallmark of severe COVID-19 is a dysregulated immune response leading to systemic inflammation causing tissue damage contributing to morbidity and mortality^7,8^. Early infection can range from asymptomatic to mild upper respiratory illness, progressing to a flu-like syndrome. Some persons then rapidly progress with acute lung injury and acute respiratory distress syndrome (ARDS)^9^. ARDS is one of the primary causes of death in COVID-19 and is characterized by inflammation-mediated damage to the alveolar-capillary barrier, resulting in reduced gas exchange and hypoxia^9,10^. Other contributors to ARDS and mortality include “cytokine storm” or cytokine release syndrome (CRS) causing a systemic inflammatory response (SIRS) accompanied by hypotension and multisystem organ failure^11,12^. However, the underlying mechanisms leading to immune dysregulation causing this cascade of events are not well understood.

Cytokines regulate numerous biological functions including the balance between innate and adaptive immune responses, and are crucial for controlling infections^2,13^. Some of the cytokines most commonly observed to be elevated in blood during severe COVID-19 include interleukin (IL)-10, IL-6, tumor necrosis factor (TNF)-ɑ, IL-1, IL-17, IL-8, and interferon (IFN)-γ^14–16^. IL-6 is a pleiotropic cytokine with a critical role in the differentiation of T and B cells and is believed to contribute to the pathogenesis of COVID-19 through inflammatory signaling^12,17^. Concentrations of circulating IL-6 are consistently reported as elevated in COVID-19 patients and have been shown to be a strong predictor of disease severity and mortality risk^2,15,18–20^. Furthermore, several clinical trials have shown that IL-6 inhibitors, such as tocilizumab and sarilumab, provide a survival benefit for some severely ill COVID-19 patients when combined with corticosteroids^21,22^, highlighting the importance of controlling systemic inflammation in severe COVID-19.

Systemic inflammation and immune activation are influenced by microbial dysbiosis and intestinal barrier damage in a range of viral infectious diseases such as influenza, respiratory syncytial virus (RSV), and HIV ^23–27^. Emerging evidence suggests that this is also true in COVID-19^28–32^. Damage to gut tissue can be mediated by direct SARS-CoV-2 infection of enterocytes or driven by systemic inflammation from a lung infection via the gut-lung axis^33,34^. Further, many studies have found that COVID-19 is associated with changes in the gut microbiome, including reduced diversity, loss of short-chain fatty acid producing commensal bacteria such as *Faecalibacterium prausnitsii*, and an enrichment of opportunistic pathogens such as *Hungatella hathewayi*, which positively correlates with pro-inflammatory cytokines such as IL-6^28–30,32,35^. In addition, we and others have demonstrated there is significant translocation of bacteria and bacterial products such as lipopolysaccharide (LPS) into the bloodstream of moderate and severe COVID-19 patients^36,37^. Circulating LPS can have many downstream consequences that induce inflammation, such as triggering the activation of monocytes and macrophages, prompting the generation of inflammatory cytokines, and may play a role in inducing ARDS^38–40^. Lipopolysaccharide-binding protein (LBP), a commonly used surrogate biomarker for LPS^41^, is a host-derived glycoprotein produced by hepatocytes and intestinal epithelial cells which binds to LPS and can enhance its inflammatory effects^38^.

Additional blood biomarkers of microbial translocation and gut barrier damage include soluble CD14 (sCD14), intestinal fatty acid binding protein (I-FABP), and zonulin, which are indicators for monocyte activation^42–45^, enterocyte damage^46^, and intestinal barrier integrity^47^, respectively. Severe COVID-19 patients have been reported to have significantly elevated concentrations of LPS, LBP, sCD14, and zonulin^31^, especially in non-survivors^48^. Understanding the connection between gut barrier integrity and systemic inflammation is an important area of investigation for understanding disease severity in COVID-19, yet this topic remains understudied. We addressed this knowledge gap by examining the differences in concentrations of circulating cytokines and markers of gut barrier function in patient plasma samples and performing a correlation analysis to understand how the relationships between these markers differ between healthy individuals and hospitalized COVID-19 patients. We then assessed the predictive potential for these biomarkers in a random forest model to distinguish between the two groups and to identify the factors most important to the model’s prediction accuracy.

## METHODS

### Study Design

In this study, we included samples from two observational cohorts at the University of California, Los Angeles (UCLA): a cohort of hospitalized patients with COVID-19 (58 individuals) and a cohort of healthy volunteers who were either hospital workers or first responders at UCLA (115 individuals). For the hospitalized patient cohort, participants were recruited from two UCLA Health hospitals in Los Angeles, CA. Inclusion criteria included hospitalization for COVID-19, age greater than 18, and confirmed positive SARS-CoV-2 RT-PCR within 72 hours of admission. Exclusion criteria included pregnancy, hemoglobin less than 8 g/dL, or inability to provide informed consent. Demographic and clinical data, including therapeutics, were collected from the electronic medical record. Clinical severity was scored using the WHO Clinical Progression Scale^49^. For this study, moderate COVID-19 included WHO scores 4-5, and severe COVID-19 included WHO scores 6-10. Initial analysis stratifying patients by severity scale did not reveal major differences between moderate and severe patients (**Supplemental Figure 1**) and thus they were combined into a single “hospitalized” grouping. Whole blood samples were collected in heparin anticoagulant tubes from each participant and plasma was separated using centrifugation and stored at -80 °C until assayed. Samples included in this analysis were collected from began in May 2020 to December 2020. Most participants provided a single plasma sample, although for 41 hospitalized patients multiple plasma samples were collected during their hospitalization, the results for which were averaged, and the mean of each participant variable was used for downstream analysis. Informed consent was collected from each participant, and the UCLA Institutional Review Board approved the study.

### Cytokine and Gut Barrier Integrity Measurements

Plasma biomarkers for gut barrier integrity were assessed using ELISAs for LBP (cellsciences, Newburyport, MA), sCD14 (R&D Systems, Minneapolis, MN), I-FABP (R&D Systems), and zonulin (MyBioSource, San Diego, CA). Plasma cytokines including IL-8, IL-17A, and IL-23 were measured using the Milliplex High Sensitivity T Cell Panel (Millipore Sigma) and IL-1ß, IL-12p70, IL-10, IL-2, IL-18, IFN-γ, TNF-α, and IL-6 were measured using a Simple Plex assay on the Ella platform (R&D Systems). All assays were performed according to manufacturer’s instructions.

### Statistical Analysis

Statistical analyses were performed in R version 4.3.3^50^. Summary statistics were generated using the *tableone* package (version 0.13.2)^51^. Any cytokine or gut barrier integrity values that were undetected were set to the assay’s limit of detection prior to analysis. Wilcoxon rank-sum tests were used to test for differences between healthy and hospitalized groups for all biomarker concentrations and corrected for multiple hypothesis testing using the Benjamini-Hochberg method^52^. Differences in age and sex between each group was determined by a Welch’s t-test and Chi-Square test respectively. For both healthy and hospitalized groups, Spearman correlations were performed between each cytokine and gut barrier integrity biomarker using the *psych* package (version 2.3.6)^53^, corrected for multiple hypothesis testing using the Benjamini-Hochberg method, and visualized using the *corrplot* package (version 0.92)^54^. The *cocor* package (version 1.1.4)^55^ was used to determine if healthy and hospitalized groups had different strengths of correlation between any two variables via a Fisher’s Z-transformation test, only testing those which had a significant adjusted Spearman correlation (p < 0.05) between either group, and correcting the output again using the Benjamini-Hochberg method. Principal component analysis (PCA) was performed using scaled and centered biomarker concentrations with the *prcomp* function from the R stats package and the *factoextra* package (version 1.07)^56^. Differences between the overall profiles of cytokine and gut barrier integrity biomarkers between groups was tested by a PERMANOVA using the adonis function from the *vegan* package (version 2.6.4)^57^ with a Euclidean distance matrix. PCA indicated a single patient (PID HOS0182) was an outlier by visual inspection and thus was excluded from the PCA to avoid spurious interpretations of the loading scores on each principal component. This patient had outlier values for IL-2, IL-12p70, and IL-1ß (an outlier was identified as above Q3 + 1.5IQR), however the participant was included in all other analyses due to the use of non-parametric tests, which limit an outlier’s influence (**Supplemental Figure 2)**. Random forest models were built using the *randomForest* package (version 4.7.1.1)^58^ to classify individuals as healthy or hospitalized using plasma concentrations of cytokines, gut barrier integrity biomarkers, or both. Each model was tuned to determine the starting number of trees and variables tried at each split then compared using area under the curve (AUC) of the receiving operating characteristic (ROC) curves using the *pROC* package (version 1.18.4)^59^. For the final model, which utilized both cytokines and gut barrier integrity biomarkers, the parameters used were 501 starting trees and mtry = 2. This model was then tested using a validation cohort as described below. The effect of age and sex on the differences between cytokines and gut barrier biomarkers was checked using multivariate linear models, using the *lmerTest* package (version 3.1-3)^60^, with log_10_ transformed values and p-value adjustment using the Benjamini-Hochberg method. Data wrangling and visualizations were produced using the *tidyverse* package (version 2.0.0)^61^ and GraphPad Prism (version 10.2).

### Validation Cohort

The random forest model uses an out-of-bag (OOB) approach with majority voting that circumvents the need to split data into testing and training sections. However, we aimed to assess the utility of the model we developed by applying it to an additional dataset of hospitalized COVID-19 patients from our laboratory^30^. Matching plasma cytokine and gut barrier integrity data was collected with the same assays from two separate studies of hospitalized COVID-19 patients, one consisting of samples from Minnesota, USA (n = 32) and another from Milan, Italy (n = 17), **Supplemental Table 1**.

## RESULTS

### Hospitalized COVID-19 patients have increased concentrations of cytokines and markers of microbial translocation

Consistent with the cytokine release syndrome described in COVID-19, the majority of cytokines measured were significantly elevated in the hospitalized patients compared to healthy volunteers including IL-1ß, IL-2, IL-6, IL-8, IL-10, IL-12p70, IL-18, IFN-γ, and TNFα (*p* < 0.05, **Figure 1**), with the largest differences being for the inflammatory cytokines IL-6 and IL-8. Biomarkers of microbial translocation including LBP and sCD14 were also significantly elevated in the hospitalized COVID-19 patients (*p* < 0.001, **Figure 1**). Furthermore, there was a significant increase in the tight junction protein zonulin in the healthy volunteers (*p* = 0.029 **Figure 1**). Dimensional reduction with PCA revealed distinct clusters for healthy and hospitalized groups using cytokines and gut barrier integrity biomarkers, and those groups were significantly distinct according to a PERMANOVA test (*p* < 0.001, r^2^ = 0.095, **Figure 2**). Most of the separation between clusters was seen on principle component 1, for which LBP, TNF-α, IL-18, IL-6, and sCD14 were the top contributing variables and directed towards the hospitalized cluster, whereas IL-17A, a homeostatic mucosal cytokine, was the top contributing factor in the direction of the healthy cluster. Of note, hospitalized patients were significantly older than the healthy volunteers (mean±sd: 56.5±13.0 *vs.* 46.0±12.2, *p* < 0.001), but there was no significant difference in sex (**Table 1**). Multivariate analysis adjusting for age and sex resulted in zonulin only trended towards significance (*p* = 0.069), though all other multivariable models were consistent with the univariate counterpart, thus age did not change the interpretation of the univariate analysis.

**Figure 1.**
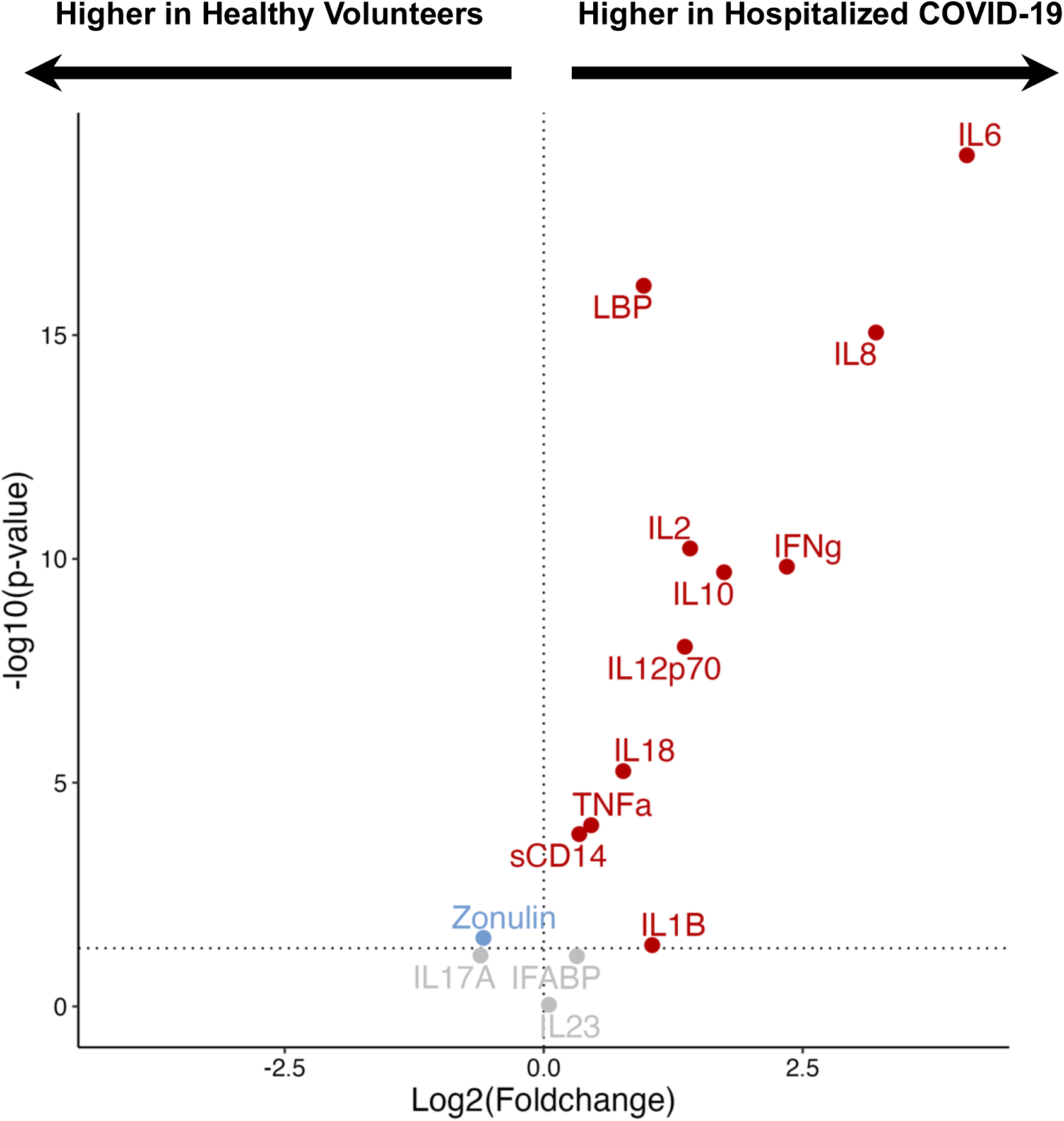
Significant differences in biomarkers between hospitalized COVID-19 patients and healthy volunteers. Volcano plot showing the log_2_ foldchange and FDR-adjusted Wilcoxon p-value for each biomarker. Red labels indicate that the biomarker was significantly higher in the hospitalized group, blue indicates significantly higher in healthy volunteers, and gray was insignificant between groups.

**Figure 2.**
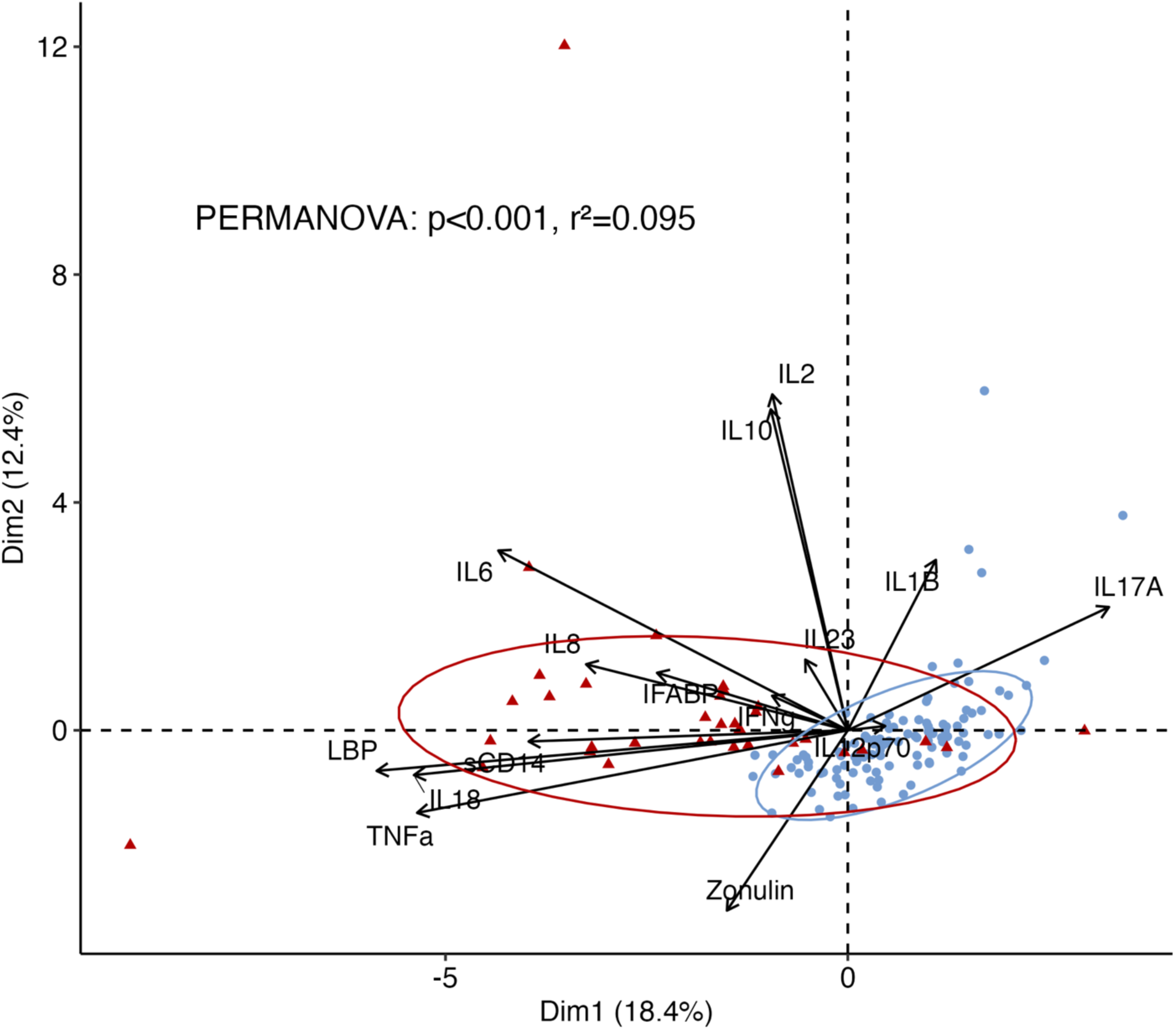
PCA of cytokines and gut barrier integrity biomarkers in healthy volunteers and hospitalized COVID-19 patients. PCA biplot showing the ordination of individuals (points) and contribution of each biomarker (arrows) across the first two principal components. Red points are hospitalized participants and blue are healthy. Individuals that are close together represent similar profiles of plasma cytokines and gut barrier integrity markers. The length of each arrow represents its contribution to the variability in the principal components and the direction indicates where that variable is best represented amongst individuals. Confidence ellipses (95%) are displayed for each group, PERMANOVA: p<0.01, r^2^ = 0.095.

**Table 1.** Demographic data of participants and plasma concentrations of biomarkers.

|  | Healthy (n=115) | Hospitalized (n=58) | p-value | Missing (%) |
| --- | --- | --- | --- | --- |
| **Age (mean (SD))** | 45.98 (12.20) | 56.53 (12.98) | <0.001<sup>a</sup> | 0 |
| **Sex (n (%))** |  |  |  |  |
| Female | 53 (46.1) | 21 (36.2) | 0.281<sup>b</sup> | 0 |
| Male | 62 (53.9) | 37 (63.8) |  |  |
| **Gut barrier markers (ng/mL)** |  |  |  |  |
| LBP | 22,848.00 (17,718.50, 33,750.00) | 50,485.33 (40,113.25, 57,018.83) | <0.001<sup>c</sup> | 0 |
| sCD14 | 1,910.97 (1,565.30, 2,339.20) | 2,421.49 (1,825.60, 3,568.11) | <0.001<sup>c</sup> | 0 |
| I-FABP | 935.60 (677.13, 1,435.67) | 1,046.90 (701.60, 1,833.21) | 0.070<sup>c</sup> | 0 |
| Zonulin | 2.33 (0.96, 5.73) | 1.59 (0.56, 2.57) | 0.022<sup>c</sup> | 1.7 |
| **Cytokines (pg/mL)** |  |  |  |  |
| IL-8 | 3.38 (1.43, 6.92) | 22.14 (10.76, 59.53) | <0.001<sup>c</sup> | 0.6 |
| IL-17A | 8.66 (2.09, 20.02) | 4.96 (3.27, 7.58) | 0.063<sup>c</sup> | 0.6 |
| IL-23 | 95.10 (29.00, 242.10) | 94.33 (35.05, 193.16) | 0.920<sup>c</sup> | 0.6 |
| IL-1β | 0.35 (0.20, 0.54) | 0.47 (0.24, 0.99) | 0.034<sup>c</sup> | 4.6 |
| IL-12p70 | 0.39 (0.39, 0.63) | 0.76 (0.58, 1.42) | <0.001<sup>c</sup> | 5.2 |
| IL-10 | 1.52 (1.03, 1.95) | 5.29 (2.41, 7.97) | <0.001<sup>c</sup> | 7.5 |
| IL-2 | 0.18 (0.18, 0.18) | 0.21 (0.18, 0.54) | <0.001<sup>c</sup> | 7.5 |
| IL-18 | 172.00 (130.00, 214.00) | 302.40 (152.49, 487.62) | <0.001<sup>c</sup> | 3.5 |
| IFN-γ | 0.58 (0.45, 0.81) | 1.87 (1.02, 11.54) | <0.001<sup>c</sup> | 3.5 |
| TNF-α | 7.58 (6.04, 9.22) | 10.44 (7.70, 13.60) | <0.001<sup>c</sup> | 4.6 |
| IL-6 | 1.36 (0.91, 2.19) | 20.27 (8.64, 76.61) | <0.001<sup>c</sup> | 4.6 |

Concentrations of biomarkers are displayed as median (IQR)
<sup>a</sup>Welch's t-test
<sup>b</sup>Chi-Square
<sup>c</sup>Wilcoxon Rank Sum Test, FDR adjusted

### Biomarker Correlations are Significantly Different Between Healthy and Hospitalized Patients

We next performed a correlation analysis between each pair of biomarkers separately for the healthy and hospitalized cohorts (**Figure 3**) and used the *cocor* R package to identify instances where the strength of the correlation between each pair of biomarkers was significantly different between the two groups (**Figure 4A**). Interestingly, this analysis revealed both shared and divergent correlation patterns between biomarkers. For example, TNFα and IL-6 were positively correlated in both healthy and hospitalized groups (r = 0.53, *p* = <0.001 and r = 0.43, *p* = 0.018, respectively, **Figure 3**). However, in the hospitalized patients, we observed positive correlations for IFABP and IL-8 (r = 0.36, *p* = 0.005), as well as LBP and IL-10 (r = 0.41, *p* = 0.005) that were significantly different from the healthy controls (r = -0.04 and r = -0.18, *cocor*: *p* = 0.028 and *p* = 0.003, respectively) (**Figure 4B-D**). Similarly, correlations between cytokines with pro-inflammatory effects often had significantly stronger positive correlations in the hospitalized patients compared to the healthy volunteers. For example, IL-6 and IL-2 were positively correlated in the hospitalized group (r = 0.47, *p* = 0.016), and significantly different from the healthy volunteers (*cocor p* = 0.011) where there was no relationship between these cytokines (r = -0.02, *p* = 0.943). Interestingly, nearly every relationship that was significant in the healthy volunteers but not in hospitalized patients was related to either IL-17A or IL-23. IL-17A was negatively correlated with LBP (r = -0.63, *p* = <0.001), zonulin (r = -0.39, *p* = <0.001), and IL-6 (r = -0.38, *p* = <0.001) in healthy volunteers and significantly different from the hospitalized group (*cocor p* < 0.05) which was not significantly correlated. Similar trends were seen with IL-23, which was negatively correlated with LBP (r = -0.61, *p* = <0.001) and IL-6 (r = -0.42, *p* = <0.001) in the healthy volunteers and significantly different from the hospitalized patients (cocor: *p* <0.001 and *p* = 0.003) where there was no relationship. This is consistent with our PCA biplot results above (Figure 2) and the biological functions of these cytokines supporting gut health, whereas during COVID-19 infection, cytokine abundance and function is significantly imbalanced.

**Figure 3.**
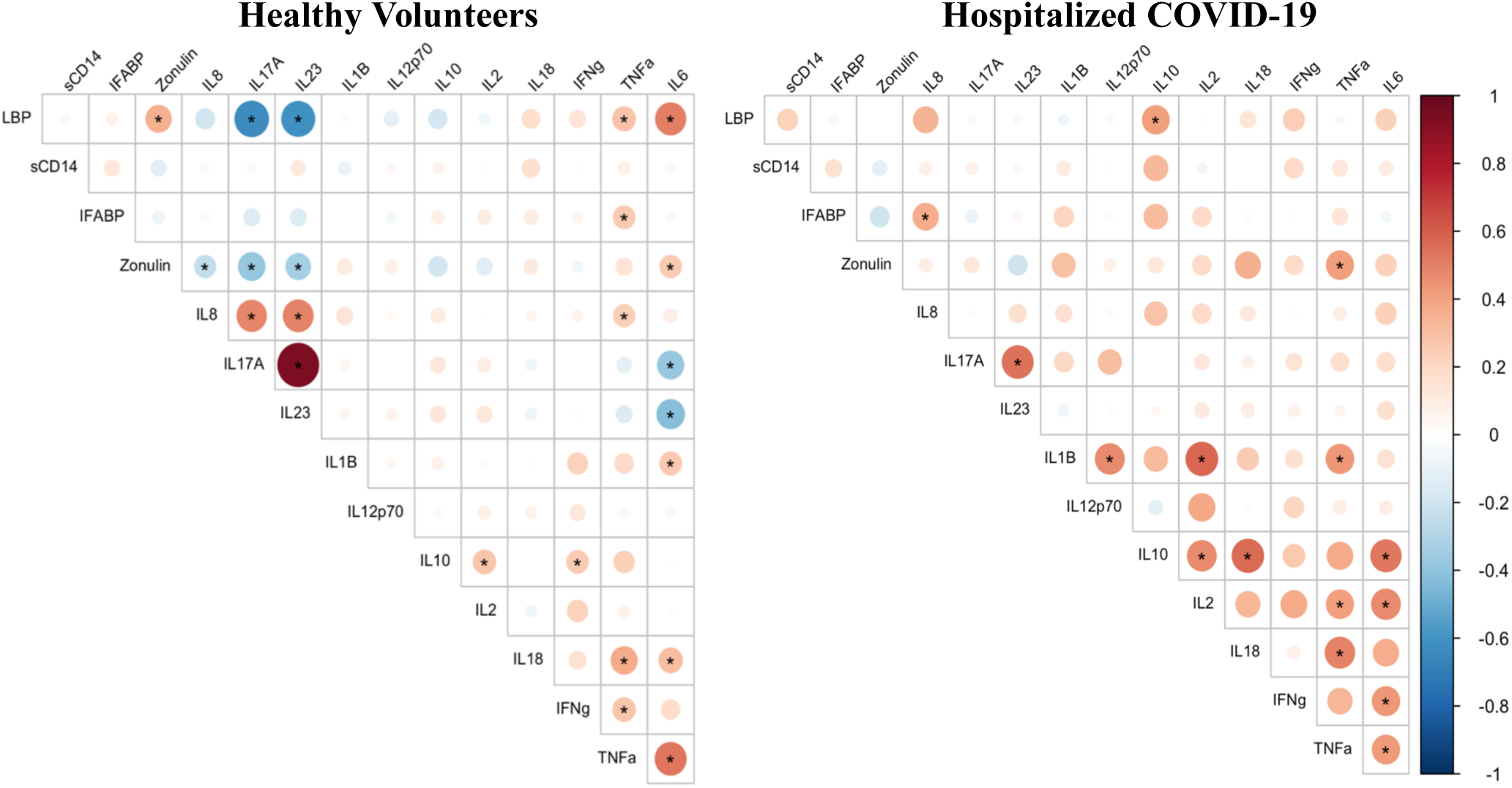
Correlations between cytokines and gut markers for hospitalized COVID-19 patients and healthy volunteers. Spearman correlations between each cytokine and gut barrier integrity biomarker, stratified by healthy volunteers and hospitalized COVID-19 patients. Asterisks denote an FDR-adjusted *p*-value <0.05. Red indicates positive correlation and blue indicates a negative correlation; the size of each circle indicates strength of the correlation.

**Figure 4.**
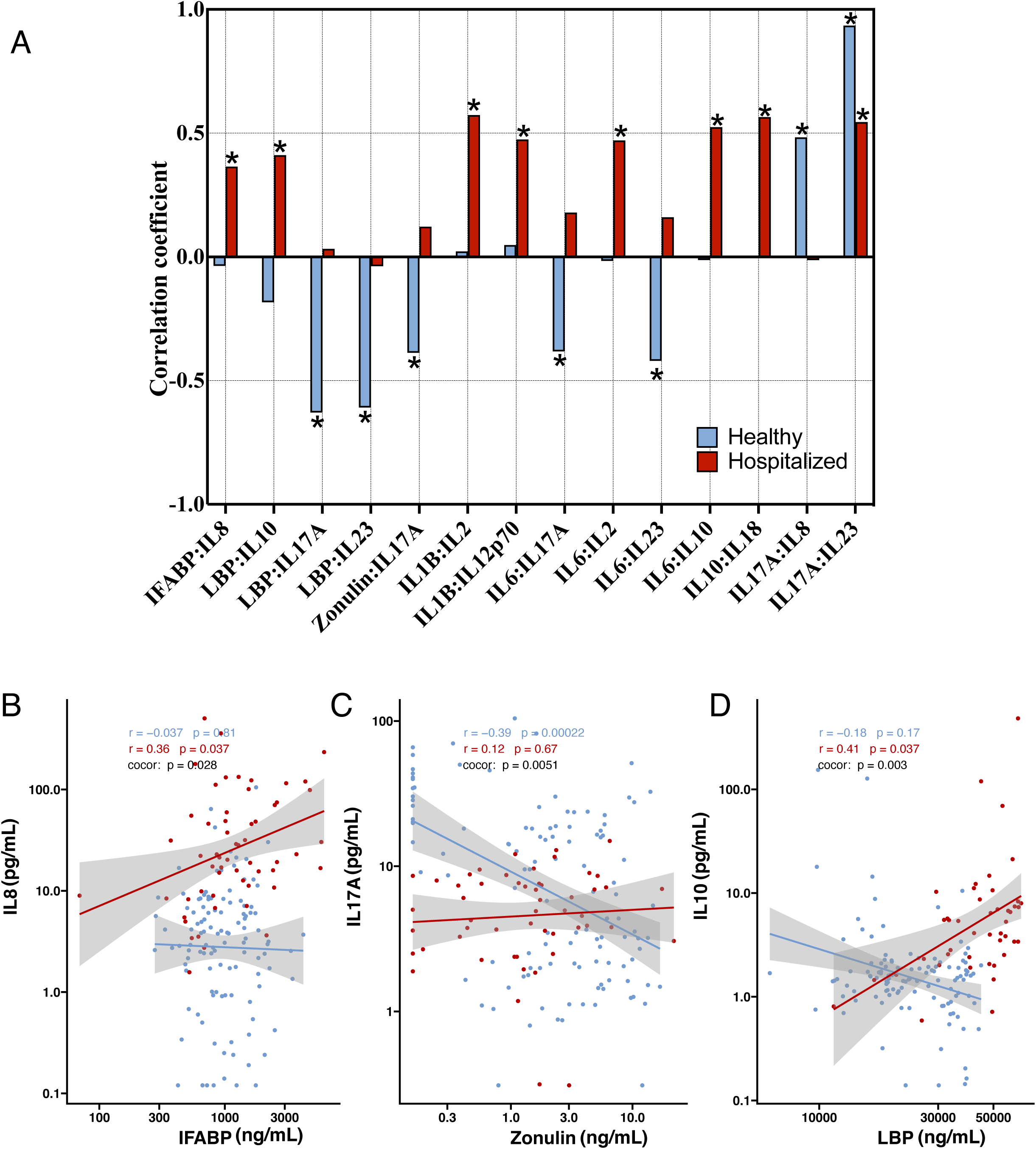
Comparison of correlation strengths between groups. Instances where strength of the correlation was significantly different between healthy volunteers and hospitalized COVID-19 patients as determined by the cocor R package (FDR *p*-value <0.05). **(A)** Each bar represents the strength and direction for a given correlation in either group, with asterisks denoting significant spearman correlations (FDR *p*-value < 0.05). **(B-D)** Representative line plots of correlations between gut barrier integrity markers and cytokines.

### Cytokine and markers of microbial translocation in plasma are predictive of hospitalized COVID-19 patients

We used supervised random forest models using cytokines, gut barrier integrity biomarkers, or a combination of both to identify predictive biomarkers of hospitalized and healthy participants. All three models performed well as measured by the area under the curve (AUC) of the receiver operator characteristic curve (ROC). The model using both cytokines and gut barrier markers had the highest AUC (97.7%), followed by cytokines only (97.2%) and then gut barrier integrity markers only (91.6%; **Figure 5A),** suggesting that these plasma biomarkers can predict hospitalized COVID-19 status. The model using both cytokines and gut barrier integrity markers was effective in classifying hospitalized and healthy individuals with a true positive percent (TPP) of 93.6%, correctly classifying 112 participants as healthy and 50 participants as hospitalized out of 173 (**Figure 5C**). Importantly, the most influential variables in model prediction, as defined by mean decrease in accuracy, were IL-6 and LBP (**Figure 5B**). This model was then tested on a validation cohort of previously unseen samples where it correctly identified 75.5% of hospitalized COVID-19 patients (n = 49; 37 out of 49, **Figure 5C**).

**Figure 5.**
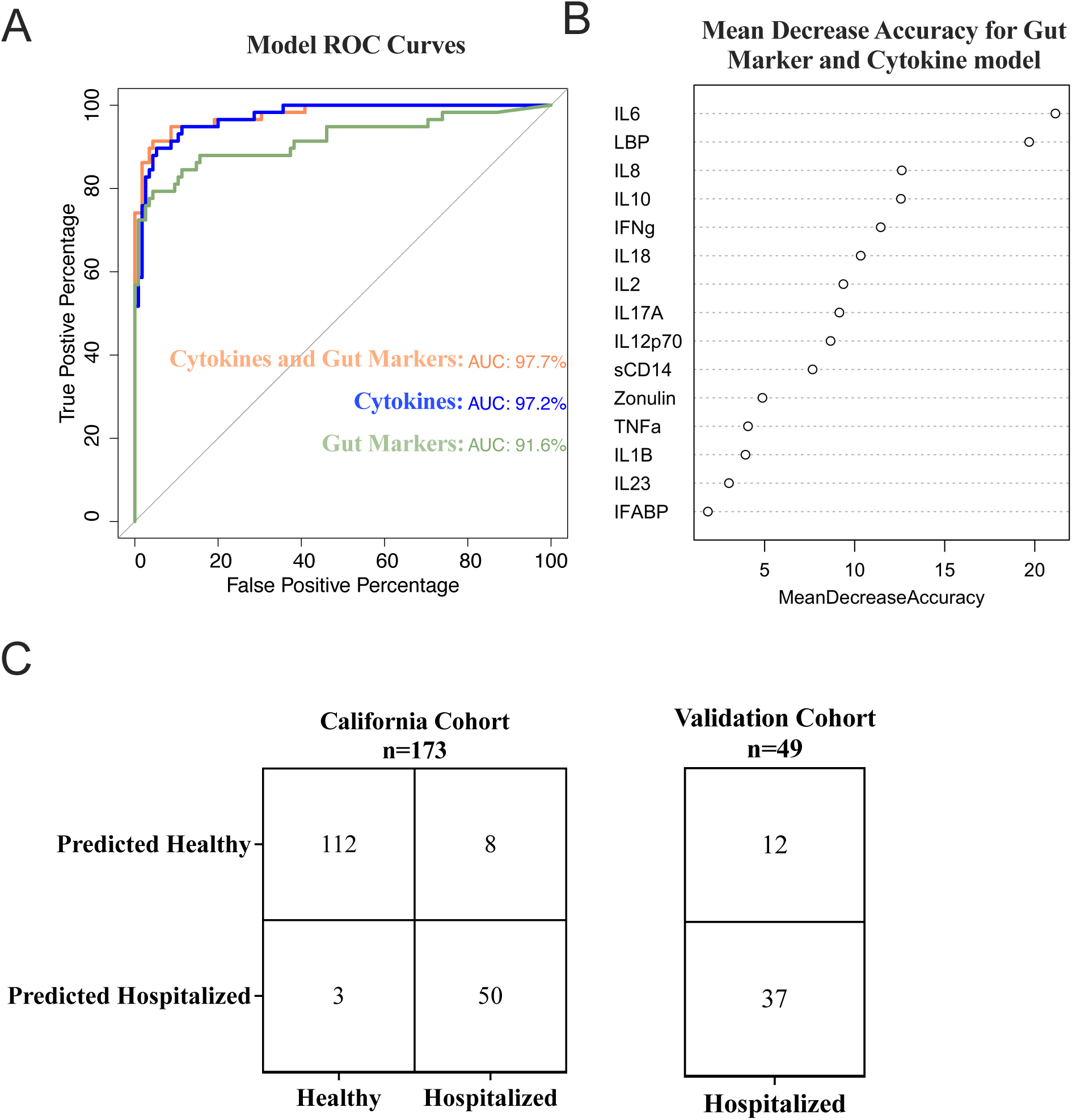
Random forest model performance. **(A)** ROC curves displaying the performance of the three different random forest models using either cytokines, gut barrier integrity markers, or both. The cytokine and gut marker model exhibited the highest area under the curve with 97.7%. **(B)** Importance plot of variable contributions to the cytokine and gut marker model accuracy. **(C)** Confusion matrices using the combined cytokine and gut marker random forest model with the original California-based cohort and the validation cohort. Each box includes the number of participants in the class the random forest model predicted compared to the group in which they are actually classified. There were no healthy controls in the validation cohort. The model correctly predicted 97.3% of individuals in the California Cohort and 75.5% in the Validation Cohort.

## DISCUSSION

COVID-19 continues to be a significant public health threat necessitating efficient early recognition of patients with an elevated risk of developing severe illness. In addition, identifying biomarkers that demonstrate differences between patients hospitalized for COVID-19 and healthy volunteers adds to the ongoing effort to understand the immune dysregulation that contributes to SARS-CoV-2 pathogenesis and that of other similar diseases. In this work, we investigated the differences in plasma concentrations of several cytokines and biomarkers of gut barrier integrity, the correlations between them, and their predictive ability in a cohort of hospitalized COVID-19 individuals and healthy controls. Our analysis is consistent with other reports showing that plasma profiles of hospitalized COVID-19 patients are characterized by elevated cytokine concentrations^14–16^. Apart from the homeostatic mucosal cytokines IL-17A and IL-23, every other cytokine we measured was elevated in the hospitalized COVID-19 patients. IL-6, IL-8, and IL-10 were some of the most elevated compared to healthy volunteers, all of which are characteristic of CRS in COVID-19^62,63^. Biomarkers of microbial translocation including LBP and sCD14 were also elevated in hospitalized patients, which suggests that translocation of microbes and microbial products from mucosal sites is increased during COVID-19 and may contribute to inflammation and disease severity.

The complex interplay between the gut microbiome, systemic inflammation, and immune dysfunction during COVID-19 is still being elucidated. Here we show that the correlative relationships between cytokines, especially those with functions related to gut barrier homeostasis such as IL-17A and IL-23, are altered in hospitalized COVID-19 patients relative to healthy controls. This change in relationship with other cytokines may indicate the normal protective effect of these cytokines in the gut is disrupted during COVID-19. IL-17A is largely produced by Th17 cells^64^ and innate lymphoid cells^65^ and under normal circumstances can have protective effects in the gut by inducing Paneth cells to upregulate antimicrobial peptides (AMPs)^66^ and regulating intestinal epithelial integrity^67^. However, dysregulation of Th17 cells and the proinflammatory activity of IL-17A has also been implicated in the pathogenesis of inflammatory disorders in the gut such as inflammatory bowel disease and Crohn’s disease^68,69^, but loss of IL-17 is associated with gut dysfunction in HIV infection^70^. Furthermore, Th17 cells are overactivated in COVID-19, which can contribute to inflammation-mediated tissue damage^71^. Although we did not see differences in the concentrations of IL-17A between groups, the changes observed in the relationships with biomarkers for microbial translocation (LBP) and tight junction proteins (zonulin) suggests that the balance of the beneficial and pathogenic effects of IL-17A may be disrupted during COVID-19, potentially contributing to disease severity. These data indicate the importance of better understanding the GI tract during COVID-19, and other infections, in the role of disease progression and overall health consequences.

A random forest model using plasma concentrations of cytokines and biomarkers of gut barrier integrity was able to accurately distinguish between healthy volunteers and hospitalized COVID-19 patients with high accuracy in this study. Previous studies corroborate these results, demonstrating that predictive models for COVID-19^72^ identify IL-6 was the most important variable in prediction accuracy in our final model. Interestingly, LBP was the second most important variable in prediction accuracy; to the best of our knowledge, few studies have used plasma LBP to predict COVID-19^73,74^ and none have shown effectiveness similar to what we observed, though it has been described as being a strong predictor of severity in community-acquired pneumonia^75^. These results provide the first evidence for LBP as a potentially novel biomarker to identify COVID-19 requiring hospitalization and should be explored further in future studies along with gut function in disease.

There are several limitations of this study. First, the time from symptom onset to hospital admittance when samples were collected was not accounted for and could influence the biomarker concentrations we measured. We also did not have a non-COVID-19 hospitalized control which may have shown if the changes observed in this study are unique to COVID-19 or present in hospitalized cases from other respiratory viral infections, stress or other factors. Furthermore, these samples were collected during 2020, which was early in the pandemic and may not fully represent the same changes that would be seen in cases with more recent SARS-CoV-2 variants, vaccinations, and after exposure or infection by previous variants. However, the data herein still provide valuable insights into disease processes that could result in novel targets for interventions for not just COVID-19 disease but other diseases.

Despite these limitations, our work here provides novel insights into the immune dysregulation and pathogenesis of COVID-19 inpatients and confirms several previously reported findings. Assessing these biomarkers and using our predictive modeling approaches will also be important to address long COVID disease severity or potentially predict long COVID. We provide strong evidence that inpatient COVID-19 is characterized by elevated pro-inflammatory cytokines and biomarkers of microbial translocation, which can be used to accurately distinguish between healthy and hospitalized individuals. We further show that the relationships between cytokines and biomarkers of gut barrier integrity are altered in COVID-19 patients requiring hospitalization, especially those that relate to gut barrier homeostasis. Similar to how monoclonal antibodies against IL-6 have been successful in treating COVID-19 by mitigating inflammatory mediated tissue damage, we suggest that interventions focused on reducing inflammation from microbial translocation and improving gut health may benefit patients with hospitalized COVID-19 patients.

## ACKNOWLEDGEMENTS

We thank all patients and the team at UCLA for taking samples during the pandemic. Measurement of cytokines and gut barrier integrity markers was performed by the Cytokine Reference Laboratory at the University of Minnesota. Funding was provided by the Department of Surgery, University of Minnesota.

## AUTHOR CONTRIBUTIONS

Conceptualization: NRK, TS, CMB, JAF, OOY Supervision: NRK, CMB, OOY, JAF

Funding acquisition: NRK, GMA, JAF, OOY

Investigation: TS, CMB, AV, KGF, GMA, NHT, JAF, OOY

Formal analysis: TS, CMB, RSC Visualization: TS, CMB

Writing – original draft: TS, CMB

Writing – review & editing: NRK, ES, CAB, RL, RSC, JAF, NHT, OOY

## DATA AVAILABILITY

All raw data and code are available upon request.

## SUPPLEMENTAL

**Supplemental Figure 1.**
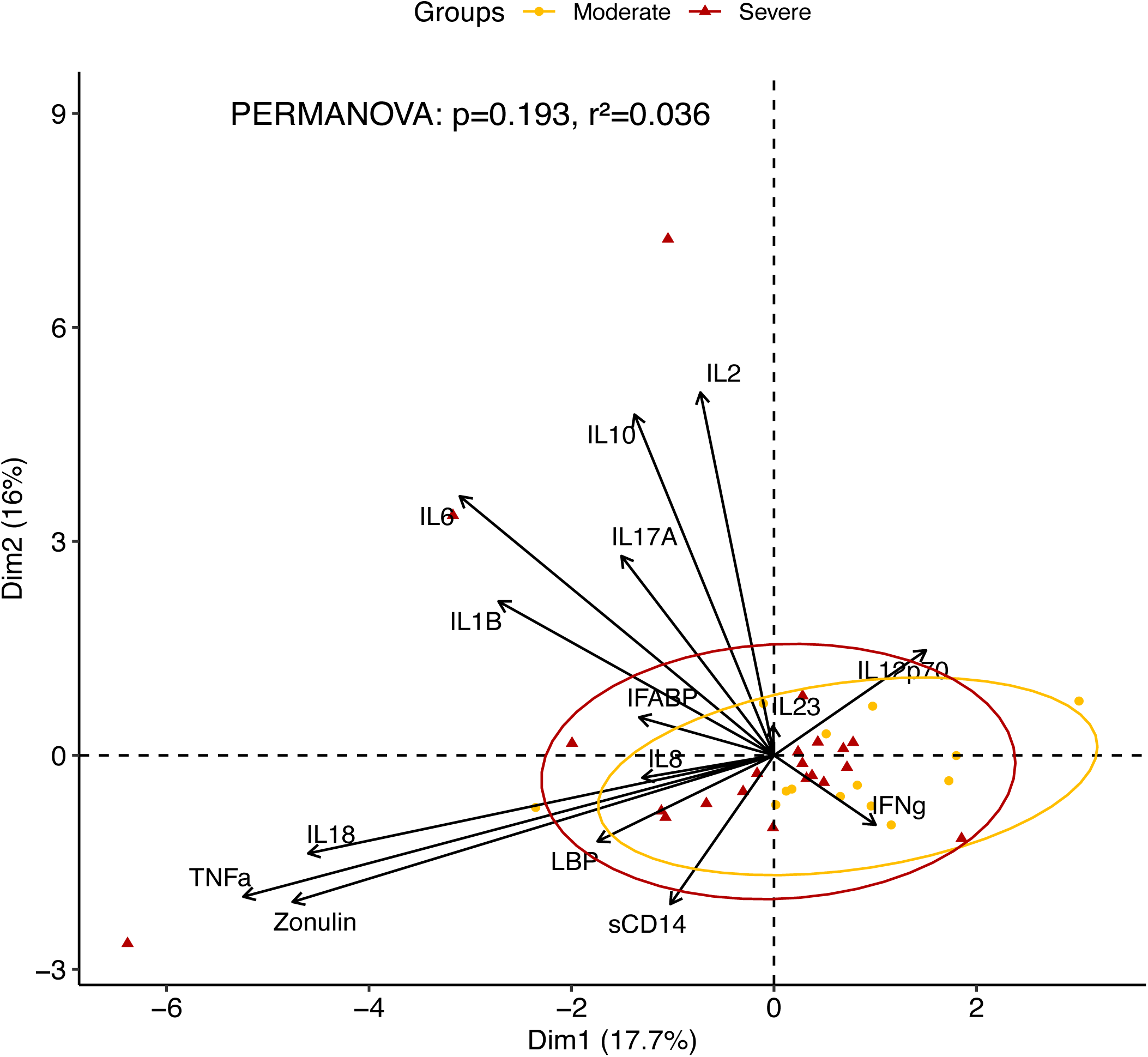
PCA of biomarkers grouped by severity. Hospitalized patients were split into their severity status as defined by the WHO scale and compared by clustering with 95% confidence intervals. Yellow ellipses and points represent Moderate COVID-19 hospitalized patients and red indicate Severe. Centroids of Moderate and Severe individuals were not significant relative to one another (PERMANOVA *p* = 0.193).

**Supplemental Figure 2.**
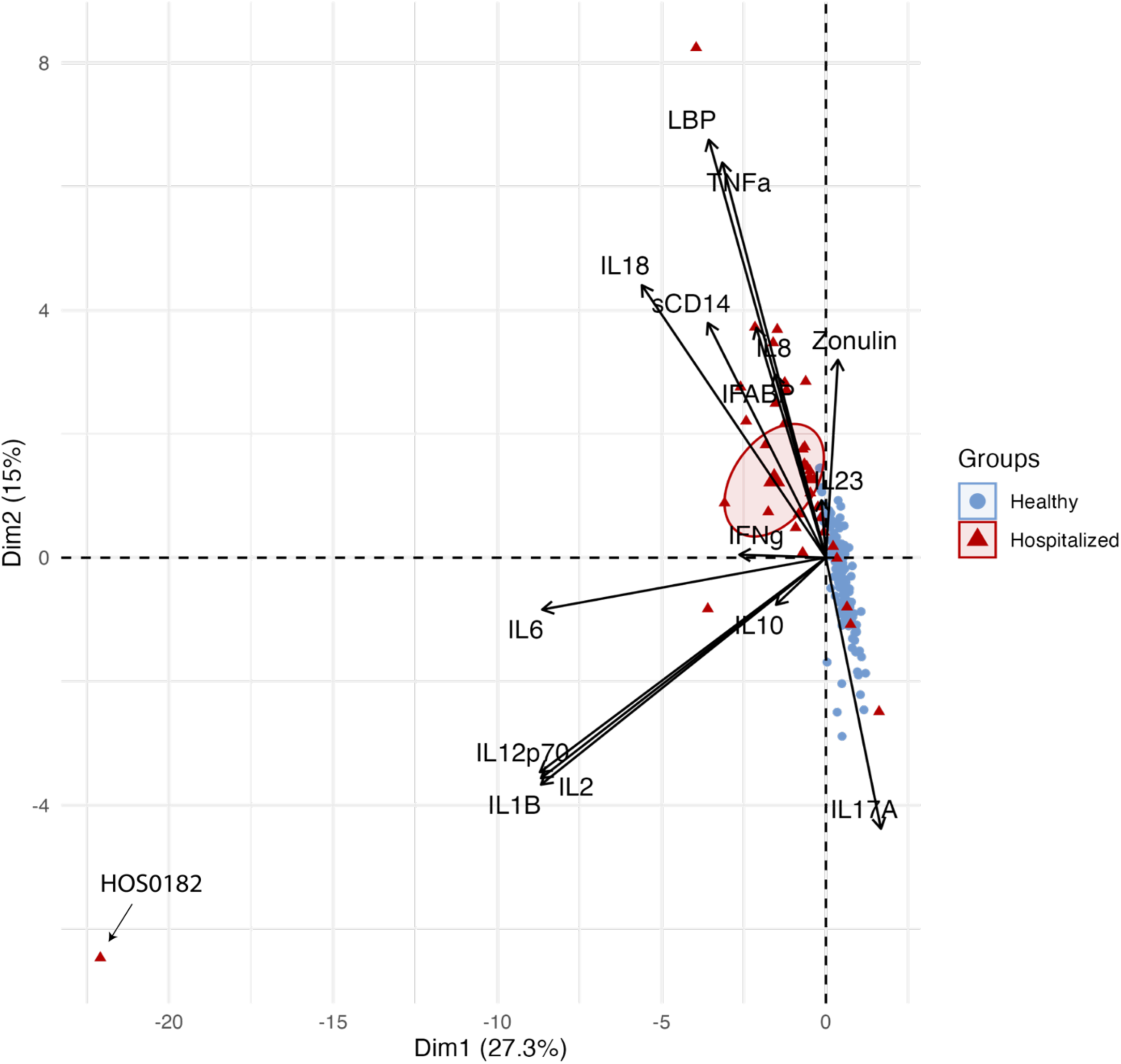
PCA of biomarkers with the outlier. Including the single individual, that is indicated as HOS0182, skews the contributors and their loadings.

**Supplemental Table 1.** Demographic data of validation cohort.

|  | **Italy (n=17)** | **United States (n=32)** | **p-value** | **Missing (%)** |
| --- | --- | --- | --- | --- |
| **Sex (n (%))** |  |  |  |  |
| Female | 2 (11.8) | 12 (37.5) | 0.089<sup>a</sup> | 4.1 |
| Male | 15 (88.2) | 18 (56.3) |  |  |

<sup>a</sup>Chi-Square

## REFERENCES

1. WHO Coronavirus (COVID-19) Dashboard https://covid19.who.int.

2. Darif, D., Hammi, I., Kihel, A., El Idrissi Saik, I., Guessous, F., and Akarid, K. (2021). The pro-inflammatory cytokines in COVID-19 pathogenesis: What goes wrong? Microb. Pathog. 153, 104799. 10.1016/j.micpath.2021.104799.

3. Liu, J., Li, Y., Liu, Q., Yao, Q., Wang, X., Zhang, H., Chen, R., Ren, L., Min, J., Deng, F., et al. (2021). SARS-CoV-2 cell tropism and multiorgan infection. Cell Discov. 7, 1–4. 10.1038/s41421-021-00249-2.

4. Groff, A., Kavanaugh, M., Ramgobin, D., McClafferty, B., Aggarwal, C.S., Golamari, R., and Jain, R. (2021). Gastrointestinal Manifestations of COVID-19: A Review of What We Know. Ochsner J. 21, 177–180. 10.31486/toj.20.0086.

5. Zeng, W., Qi, K., Ye, M., Zheng, L., Liu, X., Hu, S., Zhang, W., Tang, W., Xu, J., Yu, D., et al. (2022). Gastrointestinal symptoms are associated with severity of coronavirus disease 2019: a systematic review and meta-analysis. Eur. J. Gastroenterol. Hepatol. 34, 168–176. 10.1097/MEG.0000000000002072.

6. Bishehsari, F., Adnan, D., Deshmukh, A., Khan, S.R., Rempert, T., Dhana, K., and Mahdavinia, M. (2022). Gastrointestinal Symptoms Predict the Outcomes From COVID-19 Infection. J. Clin. Gastroenterol. 56, e145–e148. 10.1097/MCG.0000000000001513.

7. Wong, R.S.Y. (2021). Inflammation in COVID-19: from pathogenesis to treatment. Int. J. Clin. Exp. Pathol. 14, 831–844.

8. Del Valle, D.M., Kim-Schulze, S., Huang, H.-H., Beckmann, N.D., Nirenberg, S., Wang, B., Lavin, Y., Swartz, T.H., Madduri, D., Stock, A., et al. (2020). An inflammatory cytokine signature predicts COVID-19 severity and survival. Nat. Med. 26, 1636–1643. 10.1038/s41591-020-1051-9.

9. Batah, S.S., and Fabro, A.T. (2021). Pulmonary pathology of ARDS in COVID-19: A pathological review for clinicians. Respir. Med. 176, 106239. 10.1016/j.rmed.2020.106239.

10. Schifanella, L., Anderson, J., Wieking, G., Southern, P.J., Antinori, S., Galli, M., Corbellino, M., Lai, A., Klatt, N., Schacker, T.W., et al. (2023). The Defenders of the Alveolus Succumb in COVID-19 Pneumonia to SARS-CoV-2 and Necroptosis, Pyroptosis, and PANoptosis. J. Infect. Dis. 227, 1245–1254. 10.1093/infdis/jiad056.

11. Park, M.D. (2020). Macrophages: a Trojan horse in COVID-19? Nat. Rev. Immunol. 20, 351–351. 10.1038/s41577-020-0317-2.

12. Montazersaheb, S., Hosseiniyan Khatibi, S.M., Hejazi, M.S., Tarhriz, V., Farjami, A., Ghasemian Sorbeni, F., Farahzadi, R., and Ghasemnejad, T. (2022). COVID-19 infection: an overview on cytokine storm and related interventions. Virol. J. 19, 92. 10.1186/s12985-022-01814-1.

13. Kelso, A. (1998). Cytokines: Principles and prospects. Immunol. Cell Biol. 76, 300–317. 10.1046/j.1440-1711.1998.00757.x.

14. Hasanvand, A. (2022). COVID-19 and the role of cytokines in this disease. Inflammopharmacology 30, 789–798. 10.1007/s10787-022-00992-2.

15. Liu, Q.Q., Cheng, A., Wang, Y., Li, H., Hu, L., Zhao, X., Wang, T., and He, F. (2020). Cytokines and their relationship with the severity and prognosis of coronavirus disease 2019 (COVID-19): a retrospective cohort study. BMJ Open 10, e041471. 10.1136/bmjopen-2020-041471.

16. Bülow Anderberg, S., Luther, T., Berglund, M., Larsson, R., Rubertsson, S., Lipcsey, M., Larsson, A., Frithiof, R., and Hultström, M. (2021). Increased levels of plasma cytokines and correlations to organ failure and 30-day mortality in critically ill Covid-19 patients. Cytokine 138, 155389. 10.1016/j.cyto.2020.155389.

17. Scheller, J., Chalaris, A., Schmidt-Arras, D., and Rose-John, S. (2011). The pro- and anti-inflammatory properties of the cytokine interleukin-6. Biochim. Biophys. Acta BBA - Mol. Cell Res. 1813, 878–888. 10.1016/j.bbamcr.2011.01.034.

18. Udomsinprasert, W., Jittikoon, J., Sangroongruangsri, S., and Chaikledkaew, U. (2021). Circulating Levels of Interleukin-6 and Interleukin-10, But Not Tumor Necrosis Factor-Alpha, as Potential Biomarkers of Severity and Mortality for COVID-19: Systematic Review with Meta-analysis. J. Clin. Immunol. 41, 11–22. 10.1007/s10875-020-00899-z.

19. Lu, L., Zhang, H., Dauphars, D.J., and He, Y.-W. (2021). A Potential Role of Interleukin 10 in COVID-19 Pathogenesis. Trends Immunol. 42, 3–5. 10.1016/j.it.2020.10.012.

20. Santa Cruz, A., Mendes-Frias, A., Oliveira, A.I., Dias, L., Matos, A.R., Carvalho, A., Capela, C., Pedrosa, J., Castro, A.G., and Silvestre, R. (2021). Interleukin-6 Is a Biomarker for the Development of Fatal Severe Acute Respiratory Syndrome Coronavirus 2 Pneumonia. Front. Immunol. 12, 613422. 10.3389/fimmu.2021.613422.

21. RECOVERY Collaborative Group (2021). Tocilizumab in patients admitted to hospital with COVID-19 (RECOVERY): a randomised, controlled, open-label, platform trial. Lancet Lond. Engl. 397, 1637–1645. 10.1016/S0140-6736(21)00676-0.

22. REMAP-CAP Investigators, Gordon, A.C., Mouncey, P.R., Al-Beidh, F., Rowan, K.M., Nichol, A.D., Arabi, Y.M., Annane, D., Beane, A., van Bentum-Puijk, W., et al. (2021). Interleukin-6 Receptor Antagonists in Critically Ill Patients with Covid-19. N. Engl. J. Med. 384, 1491–1502. 10.1056/NEJMoa2100433.

23. Fakharian, F., Thirugnanam, S., Welsh, D.A., Kim, W.-K., Rappaport, J., Bittinger, K., and Rout, N. (2023). The Role of Gut Dysbiosis in the Loss of Intestinal Immune Cell Functions and Viral Pathogenesis. Microorganisms 11, 1849. 10.3390/microorganisms11071849.

24. Sencio, V., Machado, M.G., and Trottein, F. (2021). The lung–gut axis during viral respiratory infections: the impact of gut dysbiosis on secondary disease outcomes. Mucosal Immunol. 14, 296–304. 10.1038/s41385-020-00361-8.

25. Klatt, N.R., Canary, L.A., Sun, X., Vinton, C.L., Funderburg, N.T., Morcock, D.R., Quinones, M., Deming, C.B., Perkins, M., Hazuda, D.J., et al. (2013). Probiotic/prebiotic supplementation of antiretrovirals improves gastrointestinal immunity in SIV-infected macaques. J Clin Invest. 10.1172/JCI66227.

26. Klatt, N.R., Chomont, N., Douek, D.C., and Deeks, S.G. (2013). Immune activation and HIV persistence: implications for curative approaches to HIV infection. Immunol Rev 254, 326–342. 10.1111/imr.12065.

27. Klatt, N.R., Funderburg, N.T., and Brenchley, J.M. (2013). Microbial translocation, immune activation, and HIV disease. Trends Microbiol 21, 6–13. 10.1016/j.tim.2012.09.001.

28. Yeoh, Y.K., Zuo, T., Lui, G.C.-Y., Zhang, F., Liu, Q., Li, A.Y., Chung, A.C., Cheung, C.P., Tso, E.Y., Fung, K.S., et al. (2021). Gut microbiota composition reflects disease severity and dysfunctional immune responses in patients with COVID-19. Gut 70, 698–706. 10.1136/gutjnl-2020-323020.

29. Zuo, T., Zhang, F., Lui, G.C.Y., Yeoh, Y.K., Li, A.Y.L., Zhan, H., Wan, Y., Chung, A.C.K., Cheung, C.P., Chen, N., et al. (2020). Alterations in Gut Microbiota of Patients With COVID-19 During Time of Hospitalization. Gastroenterology 159, 944–955.e8. 10.1053/j.gastro.2020.05.048.

30. Basting, C.M., Langat, R., Broedlow, C.A., Guerrero, C.R., Bold, T.D., Bailey, M., Velez, A., Schroeder, T., Short-Miller, J., Cromarty, R., et al. (2024). SARS-CoV-2 infection is associated with intestinal permeability, systemic inflammation, and microbial dysbiosis in hospitalized patients. Microbiol. Spectr. 0, e00680–24. 10.1128/spectrum.00680-24.

31. Giron, L.B., Dweep, H., Yin, X., Wang, H., Damra, M., Goldman, A.R., Gorman, N., Palmer, C.S., Tang, H.-Y., Shaikh, M.W., et al. (2021). Plasma Markers of Disrupted Gut Permeability in Severe COVID-19 Patients. Front. Immunol. 12, 686240. 10.3389/fimmu.2021.686240.

32. Martino, C., Kellman, B.P., Sandoval, D.R., Clausen, T.M., Marotz, C.A., Song, S.J., Wandro, S., Zaramela, L.S., Salido Benítez, R.A., Zhu, Q., et al. (2020). Bacterial modification of the host glycosaminoglycan heparan sulfate modulates SARS-CoV-2 infectivity. bioRxiv, 2020.08.17.238444. 10.1101/2020.08.17.238444.

33. Dumas, A., Bernard, L., Poquet, Y., Lugo-Villarino, G., and Neyrolles, O. (2018). The role of the lung microbiota and the gut-lung axis in respiratory infectious diseases. Cell. Microbiol. 20, e12966. 10.1111/cmi.12966.

34. Lamers, M.M., Beumer, J., van der Vaart, J., Knoops, K., Puschhof, J., Breugem, T.I., Ravelli, R.B.G., Paul van Schayck, J., Mykytyn, A.Z., Duimel, H.Q., et al. (2020). SARS-CoV-2 productively infects human gut enterocytes. Science 369, 50–54. 10.1126/science.abc1669.

35. Nagata, N., Takeuchi, T., Masuoka, H., Aoki, R., Ishikane, M., Iwamoto, N., Sugiyama, M., Suda, W., Nakanishi, Y., Terada-Hirashima, J., et al. (2023). Human Gut Microbiota and Its Metabolites Impact Immune Responses in COVID-19 and Its Complications. Gastroenterology 164, 272–288. 10.1053/j.gastro.2022.09.024.

36. Sirivongrangson, P., Kulvichit, W., Payungporn, S., Pisitkun, T., Chindamporn, A., Peerapornratana, S., Pisitkun, P., Chitcharoen, S., Sawaswong, V., Worasilchai, N., et al. (2020). Endotoxemia and circulating bacteriome in severe COVID-19 patients. Intensive Care Med. Exp. 8, 72. 10.1186/s40635-020-00362-8.

37. Basting, C.M., Langat, R., Broedlow, C.A., Guerrero, C.R., Bold, T.D., Bailey, M., Velez, A., Schroeder, T., Short-Miller, J., Cromarty, R., et al. (2024). SARS-CoV-2 infection is associated with intestinal permeability, systemic inflammation, and microbial dysbiosis in hospitalized patients. Microbiol. Spectr. 12, e00680–24. 10.1128/spectrum.00680-24.

38. Gutsmann, T., Müller, M., Carroll, S.F., MacKenzie, R.C., Wiese, A., and Seydel, U. (2001). Dual Role of Lipopolysaccharide (LPS)-Binding Protein in Neutralization of LPS and Enhancement of LPS-Induced Activation of Mononuclear Cells. Infect. Immun. 69, 6942– 6950. 10.1128/IAI.69.11.6942-6950.2001.

39. Khadangi, F., Forgues, A.-S., Tremblay-Pitre, S., Dufour-Mailhot, A., Henry, C., Boucher, M., Beaulieu, M.-J., Morissette, M., Fereydoonzad, L., Brunet, D., et al. (2021). Intranasal versus intratracheal exposure to lipopolysaccharides in a murine model of acute respiratory distress syndrome. Sci. Rep. 11, 7777. 10.1038/s41598-021-87462-x.

40. Verjans, E., Kanzler, S., Ohl, K., Rieg, A.D., Ruske, N., Schippers, A., Wagner, N., Tenbrock, K., Uhlig, S., and Martin, C. (2018). Initiation of LPS-induced pulmonary dysfunction and its recovery occur independent of T cells. BMC Pulm. Med. 18, 174. 10.1186/s12890-018-0741-2.

41. Canary, L.A., Vinton, C.L., Morcock, D.R., Pierce, J.B., Estes, J.D., Brenchley, J.M., and Klatt, N.R. (2013). Rate of AIDS Progression Is Associated with Gastrointestinal Dysfunction in Simian Immunodeficiency Virus–Infected Pigtail Macaques. J. Immunol. 190, 2959–2965. 10.4049/jimmunol.1202319.

42. Lyons, J.L., Uno, H., Ancuta, P., Kamat, A., Moore, D.J., Singer, E.J., Morgello, S., and Gabuzda, D. (2011). Plasma sCD14 is a biomarker associated with impaired neurocognitive test performance in attention and learning domains in HIV infection. J Acquir Immune Defic Syndr 57, 371–379. 10.1097/QAI.0b013e3182237e54.

43. Bowman, E.R., Cameron, C.M.A., Avery, A., Gabriel, J., Kettelhut, A., Hecker, M., Sontich, C.U., Tamilselvan, B., Nichols, C.N., Richardson, B., et al. (2021). Levels of Soluble CD14 and Tumor Necrosis Factor Receptors 1 and 2 May Be Predictive of Death in Severe Coronavirus Disease 2019. J. Infect. Dis. 223, 805–810. 10.1093/infdis/jiaa744.

44. Kulkarni, M., Bowman, E., Gabriel, J., Amburgy, T., Mayne, E., Zidar, D.A., Maierhofer, C., Turner, A.N., Bazan, J.A., Koletar, S.L., et al. (2016). Altered Monocyte and Endothelial Cell Adhesion Molecule Expression Is Linked to Vascular Inflammation in Human Immunodeficiency Virus Infection. Open Forum Infect. Dis. 3, ofw224. 10.1093/ofid/ofw224.

45. Mouchati, C., Durieux, J.C., Zisis, S.N., Labbato, D., Rodgers, M.A., Ailstock, K., Reinert, B.L., Funderburg, N.T., and McComsey, G.A. (2023). Increase in gut permeability and oxidized ldl is associated with post-acute sequelae of SARS-CoV-2. Front. Immunol. 14. 10.3389/fimmu.2023.1182544.

46. Tyszko, M., Lipińska-Gediga, M., Lemańska-Perek, A., Kobylińska, K., Gozdzik, W., and Adamik, B. (2022). Intestinal Fatty Acid Binding Protein (I-FABP) as a Prognostic Marker in Critically Ill COVID-19 Patients. Pathogens 11, 1526. 10.3390/pathogens11121526.

47. Fasano, A. (2020). All disease begins in the (leaky) gut: role of zonulin-mediated gut permeability in the pathogenesis of some chronic inflammatory diseases. F1000Research 9, F1000 Faculty Rev-69. 10.12688/f1000research.20510.1.

48. Teixeira, P.C., Dorneles, G.P., Santana Filho, P.C., Da Silva, I.M., Schipper, L.L., Postiga, I.A.L., Neves, C.A.M., Rodrigues Junior, L.C., Peres, A., Souto, J.T.D., et al. (2021). Increased LPS levels coexist with systemic inflammation and result in monocyte activation in severe COVID-19 patients. Int. Immunopharmacol. 100, 108125. 10.1016/j.intimp.2021.108125.

49. A minimal common outcome measure set for COVID-19 clinical research (2020). Lancet Infect. Dis. 20, e192–e197. 10.1016/S1473-3099(20)30483-7.

50. R Core Team (2024). _R: A Language and Environment for Statistical Computing_. Version 4.3.3 (R Foundation for Statistical Computing).

51. Yoshida, K., Bartel, A., Chipman, J.J., Bohn, J., McGowan, L.Da., Barrett, M., Christensen, R.H.B., and gbouzill (2022). tableone: Create “Table 1” to Describe Baseline Characteristics with or without Propensity Score Weights. Version 0.13.2.

52. Benjamini, Y., and Hochberg, Y. (1995). Controlling the False Discovery Rate: A Practical and Powerful Approach to Multiple Testing. J. R. Stat. Soc. Ser. B Methodol. 57, 289–300.

53. Revelle, W. (2024). psych: Procedures for Psychological, Psychometric, and Personality Research. Version 2.4.3.

54. Wei, T., Simko, V., Levy, M., Xie, Y., Jin, Y., Zemla, J., Freidank, M., Cai, J., and Protivinsky, T. (2021). corrplot: Visualization of a Correlation Matrix. Version 0.92.

55. Diedenhofen, B. (2022). cocor: Comparing Correlations. Version 1.1–4.

56. Kassambara, A., and Mundt, F. (2020). factoextra: Extract and Visualize the Results of Multivariate Data Analyses. Version 1.0.7.

57. Oksanen, J., Simpson, G.L., Blanchet, F.G., Kindt, R., Legendre, P., Minchin, P.R., O’Hara, R.B., Solymos, P., Stevens, M.H.H., Szoecs, E., et al. (2022). vegan: Community Ecology Package. Version 2.6-4.

58. Cutler, F. original by L.B. and A., and Wiener, R. port by A.L. and M. (2022). randomForest: Breiman and Cutler’s Random Forests for Classification and Regression. Version 4.7-1.1.

59. Robin, X., Turck, N., Hainard, A., Tiberti, N., Lisacek, F., Sanchez, J.-C., Müller, M., code), S.S. (Fast D., Multiclass), M.D. (Hand & T., and CI), Z.B. (DeLong paired test (2023). pROC: Display and Analyze ROC Curves. Version 1.18.5.

60. Kuznetsova, A., Brockhoff, P.B., Christensen, R.H.B., and Jensen, S.P. (2020). lmerTest: Tests in Linear Mixed Effects Models. Version 3.1-3.

61. Hadley Wickham, and RStudio (2023). tidyverse: Easily Install and Load the “Tidyverse.” Version 2.0.0.

62. Shekhawat, J., Gauba, K., Gupta, S., Purohit, P., Mitra, P., Garg, M., Misra, S., Sharma, P., and Banerjee, M. (2021). Interleukin-6 Perpetrator of the COVID-19 Cytokine Storm. Indian J. Clin. Biochem. 36, 440–450. 10.1007/s12291-021-00989-8.

63. Fajgenbaum, D.C., and June, C.H. (2020). Cytokine Storm. N. Engl. J. Med. 383, 2255– 2273. 10.1056/NEJMra2026131.

64. Li, Y., Wei, C., Xu, H., Jia, J., Wei, Z., Guo, R., Jia, Y., Wu, Y., Li, Y., Qi, X., et al. (2018). The Immunoregulation of Th17 in Host against Intracellular Bacterial Infection. Mediators Inflamm. 2018, 6587296. 10.1155/2018/6587296.

65. Hueber, B., Curtis, A.D., Kroll, K., Varner, V., Jones, R., Pathak, S., Lifton, M., Van Rompay, K.K.A., De Paris, K., and Reeves, R.K. (2020). Functional Perturbation of Mucosal Group 3 Innate Lymphoid and Natural Killer Cells in Simian-Human Immunodeficiency Virus/Simian Immunodeficiency Virus-Infected Infant Rhesus Macaques. J. Virol. 94, 10.1128/jvi.01644-19. 10.1128/jvi.01644-19.

66. Brabec, T., Vobořil, M., Schierová, D., Valter, E., Šplíchalová, I., Dobeš, J., Březina, J., Dobešová, M., Aidarova, A., Jakubec, M., et al. (2023). IL-17-driven induction of Paneth cell antimicrobial functions protects the host from microbiota dysbiosis and inflammation in the ileum. Mucosal Immunol. 16, 373–385. 10.1016/j.mucimm.2023.01.005.

67. Lee, J.S., Tato, C.M., Joyce-Shaikh, B., Gulen, M.F., Cayatte, C., Chen, Y., Blumenschein, W.M., Judo, M., Ayanoglu, G., McClanahan, T.K., et al. (2015). Interleukin-23-Independent IL-17 Production Regulates Intestinal Epithelial Permeability. Immunity 43, 727–738. 10.1016/j.immuni.2015.09.003.

68. Noviello, D., Mager, R., Roda, G., Borroni, R.G., Fiorino, G., and Vetrano, S. (2021). The IL23-IL17 Immune Axis in the Treatment of Ulcerative Colitis: Successes, Defeats, and Ongoing Challenges. Front. Immunol. 12. 10.3389/fimmu.2021.611256.

69. Deng, Z., Wang, S., Wu, C., and Wang, C. (2023). IL-17 inhibitor-associated inflammatory bowel disease: A study based on literature and database analysis. Front. Pharmacol. 14. 10.3389/fphar.2023.1124628.

70. Klatt, N.R., Estes, J.D., Sun, X., Ortiz, A.M., Barber, J.S., Harris, L.D., Cervasi, B., Yokomizo, L.K., Pan, L., Vinton, C.L., et al. (2012). Loss of mucosal CD103+ DCs and IL-17+ and IL-22+ lymphocytes is associated with mucosal damage in SIV infection. Mucosal Immunol. 5, 646–657. 10.1038/mi.2012.38.

71. Martonik, D., Parfieniuk-Kowerda, A., Rogalska, M., and Flisiak, R. (2021). The Role of Th17 Response in COVID-19. Cells 10, 1550. 10.3390/cells10061550.

72. Dhar, S.K., K, V., Damodar, S., Gujar, S., and Das, M. (2021). IL-6 and IL-10 as predictors of disease severity in COVID-19 patients: results from meta-analysis and regression. Heliyon 7, e06155. 10.1016/j.heliyon.2021.e06155.

73. Rahnavard, A., Mann, B., Giri, A., Chatterjee, R., and Crandall, K.A. (2022). Metabolite, protein, and tissue dysfunction associated with COVID-19 disease severity. Sci. Rep. 12, 12204. 10.1038/s41598-022-16396-9.

74. Gomila, R.M., Martorell, G., Fraile-Ribot, P.A., Doménech-Sánchez, A., Albertí, M., Oliver, A., García-Gasalla, M., and Albertí, S. (2021). Use of Matrix-Assisted Laser Desorption Ionization Time-of-Flight Mass Spectrometry Analysis of Serum Peptidome to Classify and Predict Coronavirus Disease 2019 Severity. Open Forum Infect. Dis. 8, ofab222. 10.1093/ofid/ofab222.

75. Zobel, K., Martus, P., Pletz, M.W., Ewig, S., Prediger, M., Welte, T., Bühling, F., and CAPNETZ study group (2012). Interleukin 6, lipopolysaccharide-binding protein and interleukin 10 in the prediction of risk and etiologic patterns in patients with community-acquired pneumonia: results from the German competence network CAPNETZ. BMC Pulm. Med. 12, 6. 10.1186/1471-2466-12-6.

